# A Multi-Gene Synaptic Plasticity Array Identifies Candidate Molecular Underpinnings of Cognitive and Mood Deficits in Rats with Heart Failure

**DOI:** 10.1101/2020.08.03.234831

**Authors:** Marise B. Parent, Hildebrando Candido Ferreira-Neto, Ana Rafaela Kruemmel, Ferdinand Althammer, Atit A. Patel, Sreinick Keo, Kathryn E.Whitley, Daniel N. Cox, Javier E. Stern

## Abstract

Chronic heart failure (HF) is a serious disorder that afflicts more than 26 million patients worldwide. HF is comorbid with depression, anxiety and memory deficits that have serious implications for quality of life and self-care in patients who have HF. Despite evidence that cognitive performance is worse in HF patients with reduced ejection fraction than in HF patients with preserved cardiac function, there are few studies that have assessed the effects of severely reduced ejection fraction (≤40%) on cognition in non-human animal models. Moreover, very limited information is available regarding the effects of HF on genetic markers of synaptic plasticity in brain areas critical for memory and mood regulation. We induced HF in male rats and tested mood and anxiety (sucrose preference and elevated plus maze) and memory (spontaneous alternation and inhibitory avoidance) and measured the simultaneous expression of 84 synaptic plasticity-associated genes in dorsal (DH) and ventral hippocampus (VH), basolateral (BLA) and central amygdala (CeA,) and prefrontal cortex (PFC). We also included the hypothalamic paraventricular nucleus (PVN), which has been implicated in neurohumoral activation in HF. Our results show that rats with severely reduced ejection fraction displayed signs of polydipsia, anhedonia, increased anxiety, and impaired memory in both tasks. HF also produced a drastic downregulation of synaptic-plasticity genes in PFC and PVN, moderate decreases in DH and CeA and minimal effects in BLA and VH. Collectively, these findings identify candidate brain areas and molecular mechanisms underlying HF-induced disturbances in mood and memory.

## INTRODUCTION

Chronic heart failure (HF) is a serious cardiovascular disorder that afflicts more than 26 million patients worldwide (Ambrosy *et al.*, 2014). In the United States, it is estimated that 6.2 million people suffer from HF and the prevalence is projected to increase by 46% by 2030, more than doubling the cost of health care from $30.7 to $68.8 billion (Benjamin *et al.*, 2019). Once established, HF morbidity, prognosis and mortality are largely influenced by the degree of neurohumoral activation, which is a pathophysiological mechanism that involves increased centrally-driven sympathetic activity and elevated circulating levels of neurohormones, including vasopressin, angiotensin II and endothelins among others (Packer, 1988; Mancia, 1990; Zucker *et al.*, 2004). A major focus of research in the last decade has been to uncover the precise mechanisms underlying neurohumoral activation using experimental animal models of HF. Collectively, this research demonstrates that the hypothalamic paraventricular nucleus (PVN) is a major neuronal substrate contributing to neurohumoral activation in HF (Cohn *et al.*, 1981; Goldsmith *et al.*, 1983; Cohn *et al.*, 1984; Hodsman *et al.*, 1988), and that neuroinflammation (Kang *et al.*, 2008; Kang *et al.*, 2009; Kang *et al.*, 2010; Rana *et al.*, 2010; Yu *et al.*, 2018) and altered excitatory and inhibitory synaptic balance (Patel, 2000; Zhang *et al.*, 2002; Kleiber *et al.*, 2008; Han *et al.*, 2010; Zheng *et al.*, 2011) are likely underlying cellular mechanisms.

It is now becoming well-accepted that, in addition to neurohumoral activation and the associated systemic cardiac, renal and vascular complications, HF is comorbid with depression, anxiety and memory deficits, both in human patients (Rutledge *et al.*, 2006; Angermann *et al.*, 2012; Lossnitzer *et al.*, 2013; Cameron *et al.*, 2014; Cannon *et al.*, 2017) and experimental animal models (Toledo *et al.*, 2019; Wang *et al.*, 2019). These mood and cognitive disturbances have serious implications for quality of life and self-care in patients who have HF because hospital readmission and mortality rates are significantly higher in HF patients that have depression and/or cognitive impairments compared to HF patients without these comorbidities (Dickson *et al.*, 2007; Pressler *et al.*, 2010a; Lee *et al.*, 2013; Lossnitzer *et al.*, 2013). Thus, it is critical to identify the precise neurobiological mechanisms and substrates that underlie these mood and cognitive disturbances during HF in order to identify treatments that may improve quality of life and decrease mortality in this prevalent disease.

The hippocampus, amygdala, and prefrontal cortex (PFC) play critical roles in learning, memory and mood states. Hippocampal pathology severely impairs memory (Squire & Cave, 1991; Bird, 2017; Ekstrom & Ranganath, 2018) and produces depression and anxiety in humans (Sahay & Hen, 2007; Cole *et al.*, 2011; Kheirbek *et al.*, 2012) and non-human animals (Bannerman *et al.*, 2004; Shapiro *et al.*, 2006). Similarly, damage to the amygdala in humans and animals disrupts anxiety and impairs emotional memory (Lehmann *et al.*, 2003; Etkin & Wager, 2007; Davis *et al.*, 2010; Bocchio *et al.*, 2017). Likewise, damage to the PFC produces anhedonia, anxiety and memory impairments in non-human models (Shah & Treit, 2003; Banasr & Duman, 2008; de Visser *et al.*, 2011; Kabir *et al.*, 2017; Woloszynowska-Fraser *et al.*, 2017) and mood disorders in humans (Liu *et al.*, 2017; Belleau *et al.*, 2019; Ong *et al.*, 2019). In HF patients, neurocognitive impairments, including learning and working memory deficits, delayed recall, depression, and anxiety are correlated with significant cerebral grey matter loss and damage in brain areas relevant to cognitive function, including hippocampus, amygdala and PFC (Woo *et al.*, 2003; Almeida *et al.*, 2012; O’Donnell *et al.*, 2012; Woo *et al.*, 2015; Roy *et al.*, 2017). However, the precise mechanisms through which HF produces mood and cognitive impairments are not known. This question is difficult to address in human patients due to the influence of genetic, environmental and psychosocial variables that cannot be controlled. As a result, it is critical to develop valid animal models of mood and cognitive disturbances in HF in order to begin to identify underlying mechanisms.

Despite evidence that cognitive performance is worse in HF patients with reduced ejection fraction than in HF patients with preserved cardiac function (Jefferson *et al.*, 2011; Leto & Feola, 2014), there are limited studies that have assessed the effects of severely-reduced ejection fraction (≤40%) on cognition in non-human animal models. Moreover, very limited information is available regarding the effects of HF on genetic and molecular markers of synaptic plasticity in brain areas critical for specific types of memory (*e.g.,* spatial and emotional memory) and mood regulation. This information is necessary to begin to understand the mechanisms through which HF impairs mood and memory. As a result, in the present study we aimed to investigate the effects of HF that is characterized by severe reductions in ejection fraction on behavioral measures of mood and memory that are known to be dependent on the hippocampus, amygdala and PFC. To this end, we induced HF using a well-established ischemic HF rat model (Biancardi *et al.*, 2011; Potapenko *et al.*, 2013; Ferreira-Neto *et al.*, 2017; Ferreira-Neto & Stern, 2019) and then tested the rats in a battery of behavioral tests that are minimally stressful to assess anhedonia and anxiety (sucrose preference test [SPT] and elevated plus maze [EPM], respectively) as well as spatial working memory (spontaneous alternation; SA) and emotional long-term memory (one-trial inhibitory avoidance; IA). Moreover, we used an unbiased quantitative real time polymerase reaction (qRT-PCR) approach to identify the effects of HF on the simultaneous expression of 84 genes critical for synaptic plasticity in brain areas critical for mood and memory, including dorsal and ventral hippocampus (DH and VH, respectively), basolateral and central amygdala (BLA and CeA, respectively), PFC and PVN.

## MATERIALS AND METHODS

### Subjects

Male Wistar rats (5 wks-old, 180-200 g) were purchased from Envigo Laboratories (Indianapolis, IN) and housed in Optirat cages (Animal Care Systems, Centennial, CO, USA) on a 12 h light-dark cycle (lights on at 8:00 am) with *ad libitum* access to pelleted food and water in their home cages unless otherwise stated. Rats were housed individually following surgery and during behavioral testing. All procedures were performed in compliance with the NIH guidelines for the care of laboratory animals and approved by the Georgia State University Institutional Animal Care and Use Committee.

### Induction of heart failure (HF)

HF was induced by coronary artery ligation as described previously (Biancardi *et al.*, 2011; Ferreira-Neto *et al.*, 2017; Ferreira-Neto & Stern, 2019). Briefly, animals were anesthetized with isoflurane (2-4%) and intubated for mechanical ventilation. A left thoracotomy was performed and the heart exteriorized. The ligation was placed on the main diagonal branch of the left anterior descending coronary artery. Buprenorphine SR-LAB (0.5 mg/kg sc; Zoo Pharm, Windsor, CO, USA) was given immediately after surgery to minimize postsurgical pain. Sham control animals underwent the same procedure with the exception that the coronary artery was not occluded. Transthoracic echocardiography (Vevo 3100 systems, Visual Sonics, Toronto, ON, Canada) was performed 4 weeks after surgery under light isoflurane (3-4 %) anesthesia. The measurements of the left ventricle internal diameter and its posterior and anterior walls in systole and diastole, and the ejection fraction and fractional shortening were obtained throughout the cardiac cycle from the short-axis motion imaging mode.

### Behavioral measures of mood and memory

Behavioral testing was conducted 5-10 weeks after the induction of HF starting with the measures of mood in one cohort and memory in another. All tests were selected because they assessed mood and cognition in a minimally stressful manner and all behavioral testing and scoring were conducted by an experimenter blind to HF condition. SPT is a commonly used measure of anhedonia, which is a major symptom of depression (Willner *et al.*, 1987). SPT relies on the natural preference of rodents for sweet foods and solutions; a decreased preference for a sweet solution versus tap water is assumed to reflect depression-like behavior (Krishnan & Nestler, 2011; Liu *et al.*, 2018). Manipulations that impair DH or PFC function attenuate sucrose preference (Banasr & Duman, 2008; Liu *et al.*, 2015; Kabir *et al.*, 2017), which is consistent with evidence indicating that damage to these areas produces depression in humans (Almeida *et al.*, 2012; O’Donnell *et al.*, 2012). SPT was assessed in home cages with continuous *ad libitum* access to standard chow. Rats (Sham n = 12; HF n = 8) were first habituated to sucrose (1% wt/vol) by removing their water and giving them access to two bottles that both contained 1% sucrose for 24 h. The next day they were adapted to the preference test by giving them access to one bottle containing 1% sucrose and one containing tap water for 18 h (2 pm to 8 am). For the preference test the following day, all fluids were removed for 12 h (8 am to 8 pm) and the rats were then given access to one bottle of tap water and one bottle of 1% sucrose for 12 h. To avoid biases in drinking behavior based on the location of the bottle (i.e., left versus right), the position of the sucrose and water bottles were counterbalanced each day (Strekalova *et al.*, 2004). Sucrose preference was calculated by dividing the amount of sucrose ingested by the total amount of fluid consumed on the sucrose preference testing day. The data from four rats (Sham n = 3; HF n = 1) were excluded due to excessive leaking from bottles. At the end of the sucrose preference test the rats were weighed, and chow consumption and water intake were measured for 24 h.

The rats were then tested in the elevated plus maze (EPM) approximately 10 days later. They were handled for 3 min/day for 2 days before the EPM test and all testing was conducted during the light phase between 8:00 am and 12:00 pm. The EPM is a well-established behavioral test of anxiety (Pellow *et al.*, 1985; Kraeuter *et al.*, 2019) that relies on the natural tendency of rodents to prefer dark enclosed spaces and fear height and open spaces, and their innate motivation to explore novel environments (Montgomery, 1955; Handley & Mithani, 1984). EPM is dependent on the integrity of CeA (Ventura-Silva *et al.*, 2013), BLA (Yin *et al.*, 2019) and PFC (Lisboa *et al.*, 2010; de Visser *et al.*, 2011), which is congruent with a wealth of evidence indicating that these brain areas are involved in regulation of mood (Gallagher & Chiba, 1996; Price & Drevets, 2010; Teffer & Semendeferi, 2012; Prager *et al.*, 2016; Dixon *et al.*, 2017). The plus-shaped stainless-steel apparatus was elevated 50 cm above the floor and composed of four arms (50 cm × 12 cm). Two arms were surrounded by a 40 cm high black acrylic wall (i.e., closed arms) and two arms did not have walls (i.e., open arms). The rats were placed in the central square (12 × 12 cm) facing the closed arm, their behavior was video-recorded for 5 min, and the video-recordings were analyzed by a trained observer. The apparatus was cleaned with 70% ethanol after each rat was tested. An arm entry was defined as entering an arm with all four paws. Measures of anxiety included the percentage of time spent in the open arms [(time spent in open arm/total time spent on the maze) × 100] and the percentage of entries into the open arms [(open arm entries/ total arm entries) × 100]. Risk assessment was also used as a measure of anxiety (Holly *et al.*, 2016). Specifically, anxious animals make more stretch attend attempts, which were operationally defined as when a rat stretched its body towards an open arm after having been motionless, and then retracted (Mikics *et al.*, 2005; Holly *et al.*, 2016).

Rats (Sham n = 17; HF n = 13) were handled for 3 min/day for 2 days before SA and IA training, all testing was conducted during the light phase between 8:00 am and 12:00 pm, and the apparatuses were cleaned with 70% ethanol after each rat was tested. Rats were first assessed in SA, which is a test of spatial working memory (Richman, 1986; Lalonde, 2002). The underlying assumption is that rats must remember their visits to previous locations in order to successfully alternate between spatial locations. This is supported by the finding that removing extra-maze cues or increasing the interval between arm choices decreases alternation scores (Richman, 1986; Lalonde, 2002). SA is a task that depends on the integrity of the DH (Dillon *et al.*, 2008; Sakaguchi & Sakurai, 2020) and PFC (Delatour & Gisquet-Verrier, 1996; Woloszynowska-Fraser *et al.*, 2017). To assess SA, eat rat was placed in a Y-maze composed of three equally spaced arms (60°; 60 cm × 17.5 cm). The floor of each arm was composed of stainless steel (3.5 cm wide) and the top (14 cm wide) was covered with black Plexiglas. All of the rats were placed in the same starting arm of the Y-maze and allowed to explore the maze freely for 8 min. An alternation was defined as entering three different arms consecutively, and a percent alternation score was computed by dividing the number of alternations made by the number of arms entered minus two and then multiplying the resulting quotient by 100. The number of arms entered was used as a measure of activity.

The rats were then given training in the one-trial IA task approximately 15 days after SA testing. IA is commonly used to measure emotional, long-term memory (Jarvik & Kopp, 1967; Vazdarjanova, 2002) and is dependent on intact functioning of DH (Quevedo *et al.*, 1997; Izquierdo *et al.*, 2000; Quiroz *et al.*, 2003), VH (Giovannini *et al.*, 2005), BLA (Parent *et al.*, 1992; Nobre, 2013), CeA (Roozendaal & McGaugh, 1997), and PFC (Giovannini *et al.*, 2005; Torkaman-Boutorabi *et al.*, 2015; Torres-Garcia *et al.*, 2017). Each rat was placed into a trough-shaped apparatus (91 × 15 × 20 × 6.4 cm) that was separated into two compartments by a retractable guillotine door. The dark compartment (60 cm long) had a metal floor through which shock could be delivered. A 15-W light was placed over the lighted compartment (31 cm long) and was the only source of illumination in the room. For training, each rat was placed into the lighted compartment facing away from the retractable door and the door was then opened after the rat turned toward the door or after 12 s had passed. When the rat entered the dark compartment with all four paws, it was given a footshock (1.2 mA, 2 s) and then removed from the apparatus and returned to its home cage. The current level was assessed regularly throughout the course of the experiment with a digital multi-meter (AstroAI AM33D). For the retention test 24 h later, each rat was placed in the lighted compartment, the retractable door was lowered, and the latency (s) to cross over to the dark compartment (maximum 600 s) was recorded and used as a measure of retention. Footshock was not delivered on the retention test.

### Unbiased qRT-PCR array approach to assess the impact of HF on 84 molecular markers of synaptic plasticity

Approximately 10 weeks after surgery, rats (Sham n = 5; HF n = 5) were anesthetized deeply with pentobarbital (Euthasol, Virbac, ANADA #200-071, Fort Worth, TX, USA, 390 mg/kg, i.p.), decapitated quickly with a guillotine, and their brains dissected, frozen in 2-methylbutane (Fisher Scientific, USA) at −40°C and then stored at −80°C. To collect micropunches, brains were allowed to reach −20°C and then mounted in a cryostat (CM3050S; Leica Microsystems, Wetzlar, Germany) and brain slices (300 μm) were obtained (infralimbic and prelimbic PFC, 3.7 to 2.2 mm; PVN, −1.4 to −2.12 mm; BLA and CeA, −1.8 to −3.14 mm; DH, −2.8 to −4.16 mm and VH, −4.8 to −6.04 mm from bregma) (Paxinos & Watson, 2006). A 2.0 mm stainless-steel punch was used to collect four central punches/rat from PFC, PVN, and DH and 10 bilateral punches/rat from VH. A 1.0 mm punch was used to collect eight bilateral punches from BLA and CeA. The micropunched tissues were kept on dry ice until RNA extraction. RNA extraction and isolation were performed using the miRNAeasy Mini kit (Qiagen, Hilden, Germany) and the QIAzol Lysis Reagent (Qiagen) following manufacturers’ protocols. RNA concentration was measured using NanoDrop One (Thermo Scientific, Waltham, Massachusetts, USA) and was in the range of 115 – 220 ng/µL prior to cDNA synthesis. cDNA synthesis was performed using the RT2 First Strand Kit (Cat. No. 330401, Qiagen) and the SimpliAmp Thermal Cycler (Applied Biosystems, Thermo Fisher Scientific) following the manufacturers’ protocols. The gene array reaction was performed using the RT^2^ Profiler™ Rat Synaptic Plasticity PCR Array (Cat. No. 330231, Qiagen) following the manufacturer’s protocol, and all reactions were performed in triplicate for each brain region and treatment condition using a Roche Lightcycler 96 (Roche Life Science, Indianapolis, IN, USA). This kit measures the expression of 84 genes critical for synaptic alterations implicated in memory and mood, including immediate early genes (IEG)s, genes implicated in long-term potentiation (LTP) and long-term depression (LTD), cell adhesion and extracellular matrix molecules, CREB cofactors, and neuronal receptors (plus five housekeeping genes and two negative controls).

### Validation of gene array results

To validate the array findings, qRT-PCR was conducted using specific 10x QuantiTect primers diluted in 1.1 mL TE pH 8.0 (Qiagen). The genes that were analyzed were *Adcy1* (QT00386421), *Arc* (QT00373086), *Homer1* (QT00391111), *Mapk1* (QT00190379), and *Nr4a1* (QT00181048). *B2m* (QT00176295) was used as the reference gene. All individual qRT-PCR reactions (brain region, primer and condition) were performed in triplicate.

### Statistical analyses

Statistical analyses were performed using IBM SPSS Statistics 25 (IBM, Armonk, New York) and GraphPad Prism 7 (GraphPad Software, San Diego, CA). Data points that were two standard deviations from the mean were considered outliers and excluded from further analyses. Given that we predicted that HF would impair mood and memory and increase thirst (Allida *et al.*, 2015), unpaired one-tailed *t-tests* were used to assess differences between Sham and HF rats in the behavioral tests and measures of water intake. Two-tailed tests were used to compare chow intake and body mass. Behavioral and body mass data are expressed as means ± standard error of the mean (SEM), and an alpha level of 0.05 was used as the criterion for statistical significance. Echocardiographic parameters between sham and HF rats were compared using unpaired *t-tests*.

We used the Multi Ontology Enrichment Tool (MOET) to explore functional categories of genes that were altered in HF within specific brain regions. MOET provides an unbiased analysis of groups of genes within each brain region that are functionally related, co-occur and are statistically enriched in a given gene population. To identify functional classes of genes that are statistically over-represented amongst the genes that are differentially expressed during HF, MOET uses the Gene Ontology (GO) consortium, which provides detailed annotations of genes with respect to their known or putative molecular function, cellular localization and biological processes (Smith *et al.*, 2020). MOET provides a quantitative measurement of enrichment based on well accepted statistical methods including hypergeometric distributions to calculate p-values and correct the p-values with Holm-Bonferroni corrections and presents the number of genes involved in different annotations and the corresponding significance level in disease, pathway, biological, cellular and molecular ontologies. Four numbers are used to calculate each p-value

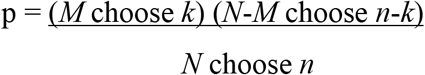

where *n* is the number of genes in our list, *N* is the number of genes in the reference population, *k* is the number of genes annotated with this item in our list and *M* is the number of genes annotated with items in the reference population. The p-value was considered significant when p<0.01. The data that support the findings of this study are available from the corresponding author upon reasonable request.

## RESULTS

### HF rats had severely reduced ejection fraction

Figure 1 shows representative echocardiographic images from a Sham and HF rat and the mean heart function values obtained are shown in Table 1. Compared to Sham, HF rats showed a significant increase in the left ventricle diameter and volume during systole and diastole (p<0.0001) and a significant reduction in the left ventricle posterior and anterior thickness during systole (p<0.0001). Also, the HF rats exhibited a significant reduction in the percentage of fractional shortening and ejection fraction (p<0.0001).

**Table 1.** Summary data of echocardiography measurements of left ventricular parameters obtained from Sham and HF rats. Values are expressed as means ± SEM; Sham n = 21; HF n = LVID, d and s: left ventricle internal dimension during diastole and systole; LVAW, d and s: left ventricle anterior wall thickness during diastole and systole; LVPW, d and s: left ventricle posterior wall thickness during diastole and systole; LV Vol, d and s: left ventricle volume during diastole and systole; EF, ejection fraction; FS, fractional shortening. ****p<0.0001 vs. Sham.

|  | Sham | HF |
| --- | --- | --- |
| LVID,d (mm) | 7.98 ± 0.18 | 10.68 ± 0.20**** |
| LVPW,d (mm) | 2.13 ± 0.11 | 2.02 ± 0.17 |
| LVAW,d (mm) | 1.97 ± 0.05 | 2.02 ± 0.12 |
| LV Vol,d (μL) | 347.34 ± 17.05 | 659.89 ± 27.97**** |
| LVID,s (mm) | 3.84 ± 0.17 | 9.19 ± 0.21**** |
| LVPW,s (mm) | 3.41 ± 0.13 | 2.18 ± 0.16**** |
| LVAW,s (mm) | 3.49 ± 0.08 | 2.62 ± 0.15**** |
| LV Vol,s (μL) | 68.35 ± 6.69 | 476.95 ± 24.40**** |
| EF % | 81.02 ± 1.39 | 28.08 ± 1.55**** |
| FS % | 52.19 ± 1.50 | 14.08 ± 0.84**** |

**Figure 1.**
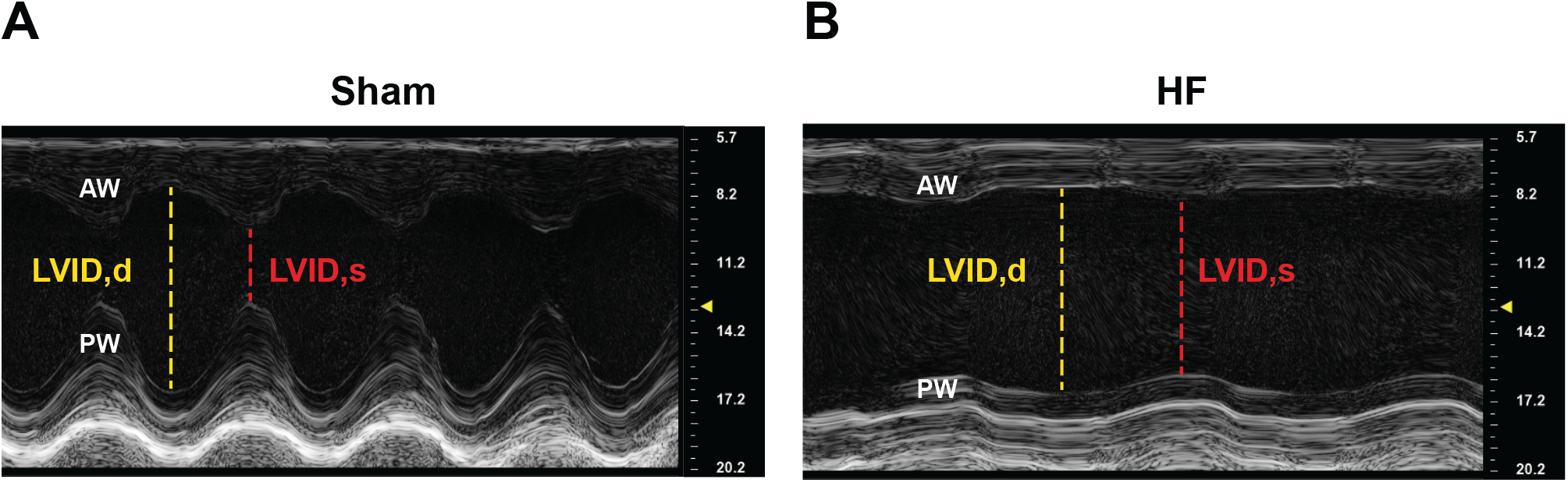
HF rats displayed severely reduced left ventricular function. Representative echocardiograph images of the movement-mode, short-axis view from a Sham **(A)** and a Heart Failure (HF) rat **(B)**. AW, anterior wall; PW, posterior wall; LV, left ventricle; LVID, d/s: left ventricle internal diameter in diastole and systole.

### HF rats displayed signs of polydipsia, anhedonia and increased anxiety

Our results show that HF rats consumed more sucrose during the first sucrose exposure day when they were only given access to sucrose (p = 0.018; Fig. 2A, n= 9 and 7 for Sham and HF, respectively). More importantly, HF rats had a lower preference for sucrose on the preference test (p = 0.05; Fig. 2B) than Sham rats. When the rats were weighed and chow consumption and water intake were measured for 24 h after the SPT, HF rats consumed more water (p = 0.02; Figure 2C) than Sham rats, but had comparable chow intake (p>0.05; Fig. 2D) and body mass (p>0.05; Fig. 2E). Together, these studies indicate that HF rats ingested more fluids than Sham rats and that HF rats also had a significantly decreased preference for sucrose, which is a sign of anhedonia (Krishnan & Nestler, 2011; Liu *et al.*, 2018).

**Figure 2.**
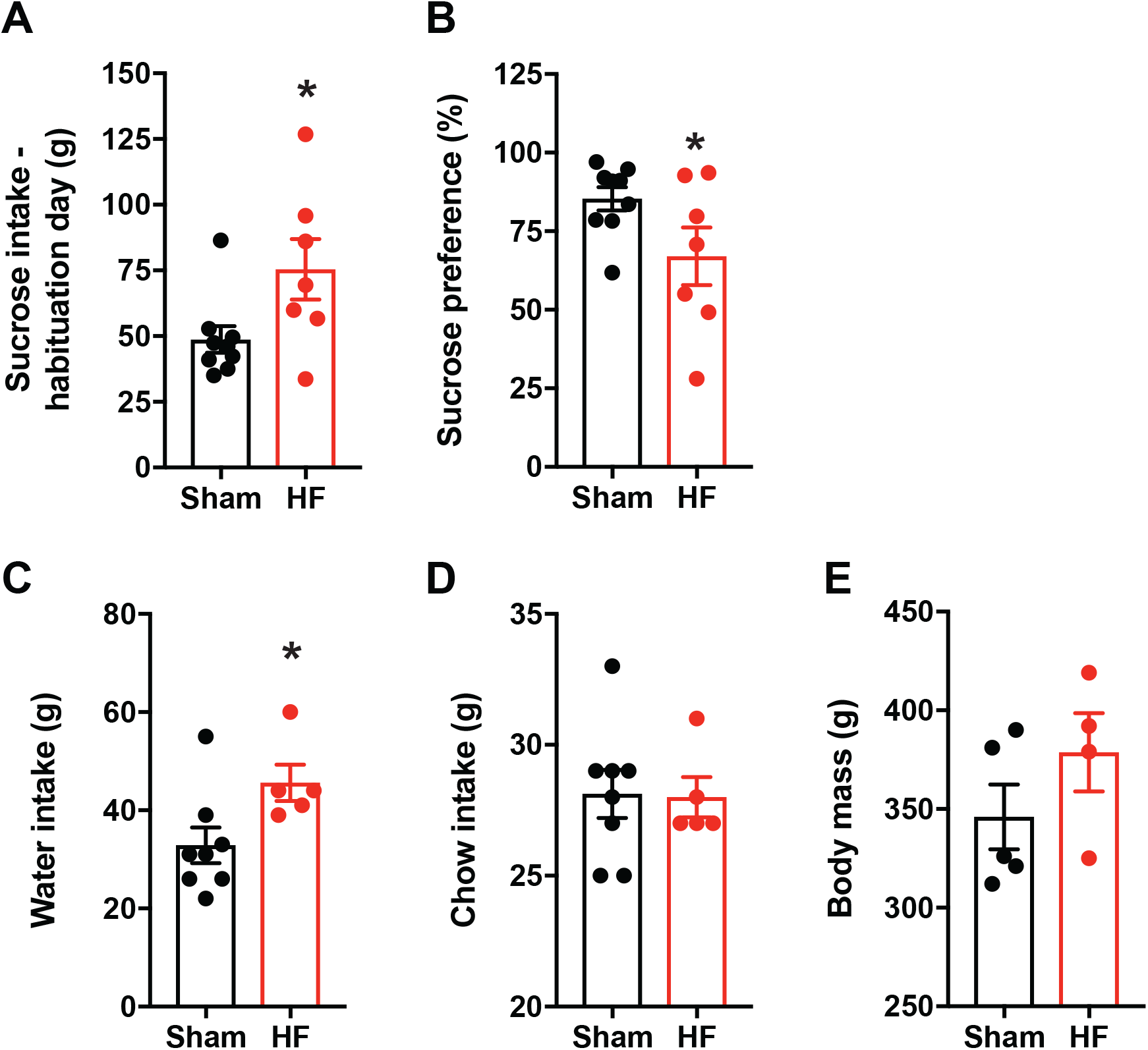
HF produced polydipsia and anhedonia. **(A)** In the sucrose preference test, HF rats consumed more sucrose than Sham controls during the first exposure day when they were only offered sucrose, but more importantly had a **(B)** decreased preference for sucrose on the sucrose preference testing day. **(C)** During a subsequent 24 h period, HF rats consumed more water than Sham rats, but had comparable chow intake **(D)** and body mass **(E)**. *p<0.05 vs. Sham.

HF rats displayed several signs of anxiety in the EPM test (n= 12 and 8 for Sham and HF, respectively). Specifically, HF rats spent less time in the open arms of the maze (p = 0.048; Fig. 3A,), tended to make fewer entries into the open arms (p = 0.057; Fig. 3B) and made more stretch attend attempts than Sham rats (p = 0.041; Fig. 3C).

**Figure 3.**
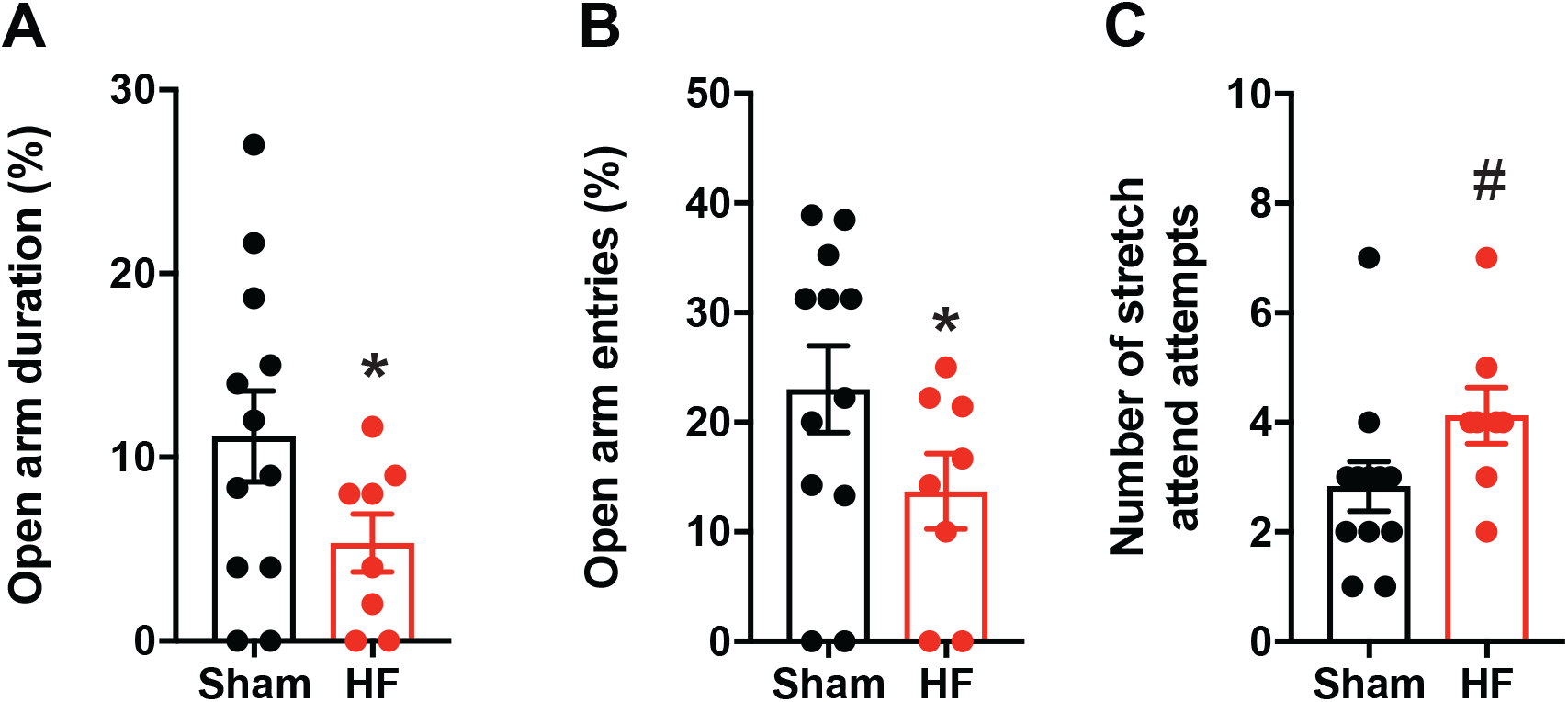
HF increases anxiety and risk assessment. **(A)** In the EPM, HF rats spent significantly less per cent time in the open arms, **(B)** tended to make fewer per cent entries into the open arms and **(C)** made more stretch attend attempts. *p<0.05 and #p = 0.057 vs. Sham.

### HF rats had impaired spatial working memory and emotional, long-term memory

In the SA task (n= 17 and 13 for Sham and HF, respectively), HF rats made fewer alternations between arms (p = 0.026; Fig. 4A), indicative of impaired spatial working memory. Although HF significantly disrupted the sequence of arm entries (i.e., percent alternation), it did not affect the number of arms the rats explored in the maze (p>0.05; Fig. 4B), suggesting that impaired alternation was not due to changes in activity levels.

**Figure 4.**
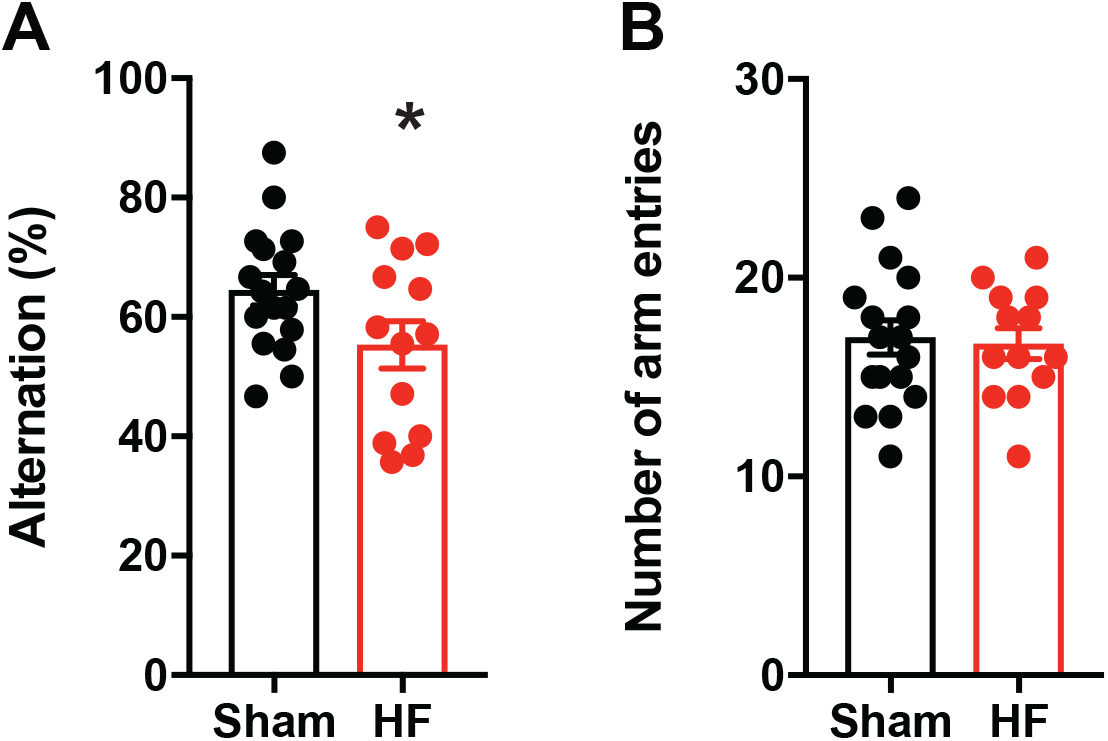
HF impairs spatial working memory without affecting activity levels. **(A)** In the SA test, HF rats **made** significantly fewer alternations between arms, but **(B)** entered a comparable number of arms. *p<0.05 vs. Sham.

On the IA retention test (n= 17 and 12 for Sham and HF, respectively), HF rats displayed significantly shorter latencies to enter the compartment in which they had previously received shock than Sham rats (p = 0.025; Fig. 5A), a result that is indicative of impaired emotional, long-term memory. Of note, rats with HF did not have shorter latencies than Sham rats on the training day (p>0.05; Fig. 5B), suggesting that the HF-induced decreases in the retention latencies were not due to hyperactivity.

**Figure 5.**
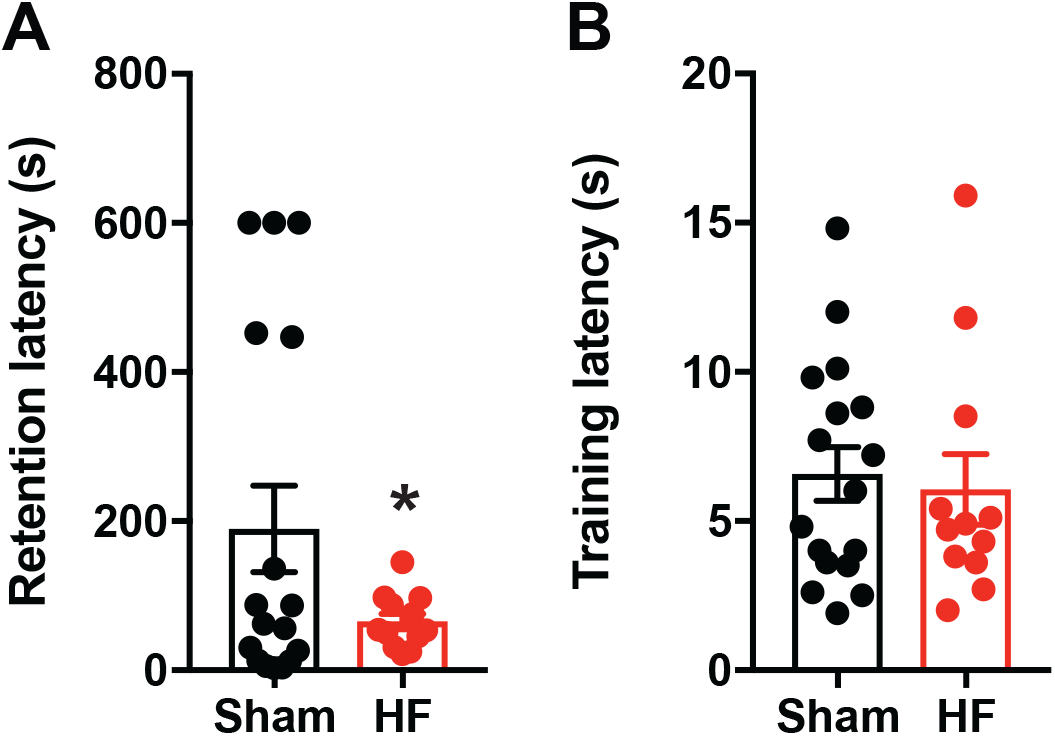
HF impairs emotional, long-term memory. **(A)** In the IA task, HF rats entered the dark/shock compartment more quickly than Sham rats on the retention test, **(B)** but their latencies did not differ during training. *p<0.05 vs. Sham.

### HF drastically downregulates the expression of several genes involved in synaptic plasticity in the PFC, PVN, CeA and DH

To begin to identify potential molecular mechanisms underlying the HF-induced changes in mood and memory that we observed, we determined whether HF impacted expression of genes involved in synaptic plasticity in brain regions known to be critical neuronal substrates for mood and memory (i.e., PFC, DH, VH, BLA, and CeA). We also included the PVN, a region that is well-established to play a major role in neurohumoral activation in this disease state, in which altered synaptic function and plasticity were described previously (Han *et al.*, 2010; Potapenko *et al.*, 2011; Potapenko *et al.*, 2012, 2013; Stern & Potapenko, 2013).

Given the extensive number of genes sampled in the array, and in order to prioritize for further validation and interpretation those genes that were more robustly affected, we set a qualitative and arbitrary threshold of a +/− 40% change in expression. Using this criterion, we generated a heatmap graph (Figure 6A) that exhibits the profile of genes whose expression were robustly upregulated (≥ +40% change, in blue range color), downregulated (≤ +40% change, in red range color), or showed no changes (white rectangles) in expression when comparing HF to sham rats across the six different brain regions. Figure 6B provides a summary of the main changes observed by brain region. Overall, the results showed a drastic HF-induced downregulation of synaptic plasticity-related genes compared to Sham rats, with ~ 87% of genes showing a downregulation (≤ +40% in expression) in at least one brain region. Conversely, increased expression (≥ +40% in expression) was observed only in ~5% of genes tested. The PFC and PVN exhibited the largest number of down regulated genes during HF, with ~80% and ~54% genes decreased, respectively. A milder change was observed in the DH and CeA, where HF downregulated ~20% of genes tested compared to Sham rats. Lastly, HF had minimal effects on synaptic plasticity genes in the BLA and VH, with less than 10% of genes being altered (Fig. 6B). Interestingly, the few genes that showed increases were mostly (3/4 genes, 75%) restricted to the VH.

**Figure 6.**
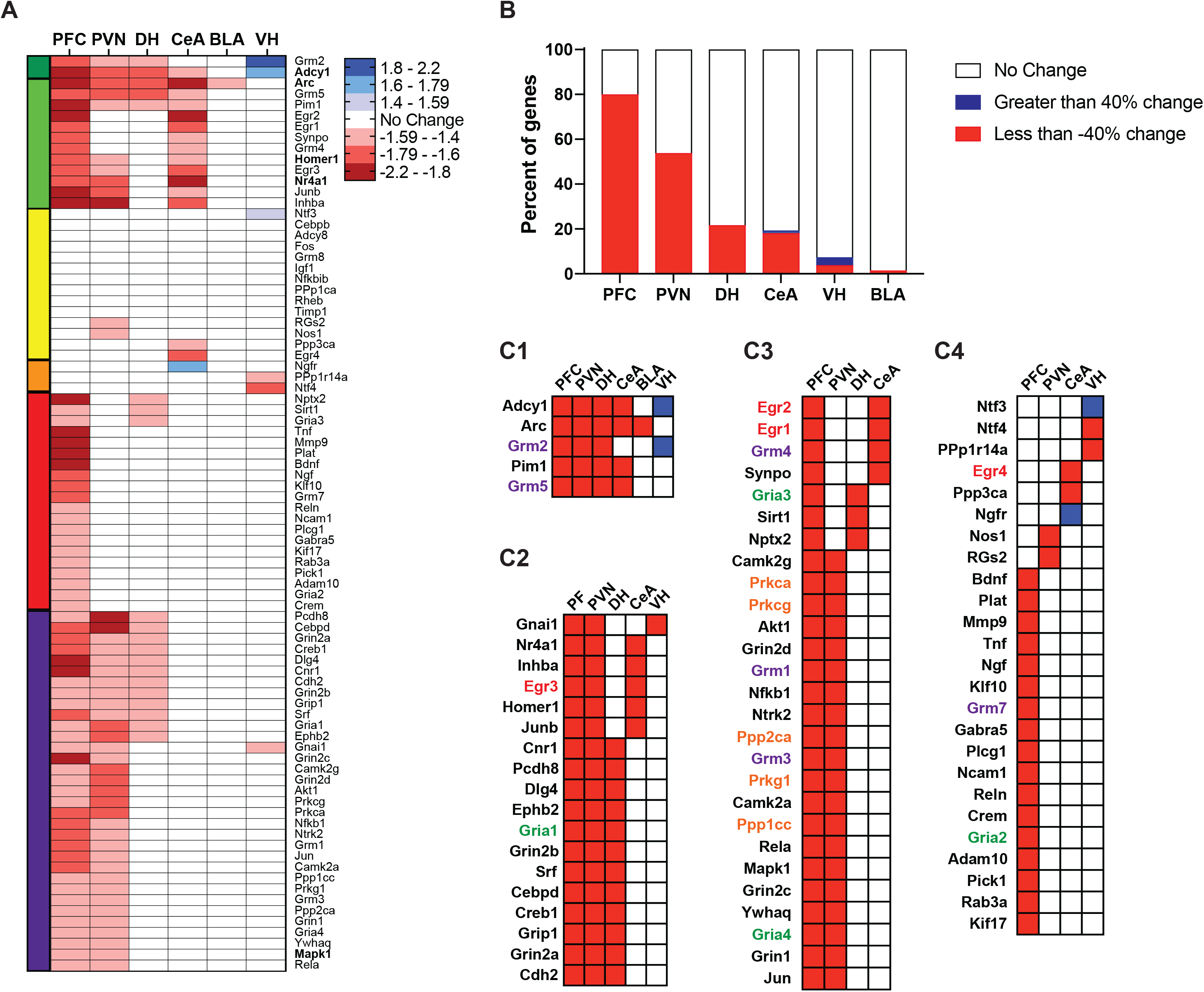
HF significantly alters synaptic plasticity-related gene expression. (**A**) Heatmap graph showing the level of expression of different genes regulated by HF in relation to sham rats’ gene expression. Downregulation below 40% change is shown in the red range, upregulation above 40% change is shown in the blue range and the expression of genes that did not change below or above 40% is shown in white. *Adcy1*, *Arc*, *Homer1*, *Mapk1* and *Nr4a1*, the genes used for the qPCR validation, are in bold. (**B**) Bar graph showing the percentage of genes that were upregulated (blue), downregulated (red), or did not change (white) during HF in PFC, PVN, DH, CeA, BLA and VH. (**C1-C4**) Heatmap graphs depicting clusters of genes that were upregulated (blue), downregulated (red) or not affected (white) due to heart failure. Clusters are organized according to genes that changed with HF vs. Sham in four to five brain areas (**C1**); three brain areas (**C2**); two brain areas (**C3**) or only one brain area (**C4**). Categories written in: red represents *Egr* family; green represents *Gria* family; purple represents *Grm* family; orange represents kinases and phosphatases related genes.

HF downregulated several important groups of genes, including IEGs (activity-regulated cytoskeletal-associated protein [*Arc*; aka Arg3.1], the *Egr* family of zinc fingered transcription factors, and *Jun*), genes encoding for glutamate receptors, predominantly metabotropic receptors (*Grm1, −2, −3, −4, −5, −7*), but also ionotropic NMDA receptor subunits (*Grin2a* and *Grin2b*). HF also downregulated various types of protein phosphatases (*Ppp1cc* and *Ppp2ca*) and protein kinases (*Prkca, Prkcg* and *Prkg1*) in the PFC and PVN. Figure 6C depicts genes grouped according to the number of brain areas in which they were impacted, with genes changing in four to five brain areas grouped in Cluster 1 (C1), three in Cluster 2 (C2), two in Cluster 3 (C3) and only one brain area in Cluster 4 (C4). The genes whose expression changed in most of the brain regions tested during HF included *Arc*, *Adcy1* (type 1 adenylyl cyclase), *Grm2* and *Grm5*, and the proto-proto-oncogene serine/threonine-protein kinase Pim-1 (*Pim1;* see C1). Interestingly, another gene directly associated to metabotropic glutamate receptor (mGluR) function and synaptic plasticity, *Homer1* (Fig. 6A, light green cluster), was also downregulated during HF in the PFC, PVN and CeA. Cyclic adenosine monophosphate (cAMP) responsive element binding protein 1 gene (*Creb1*), a gene critically linked to vascular dementia, learning, memory and cognitive abilities (Brightwell *et al.*, 2004; Sadamoto *et al.*, 2010; Han *et al.*, 2018), was also reduced by HF in the PFC, PVN and DH (Fig. 6A, purple cluster).

We then used the Multi Ontology Enrichment Tool (MOET), which provides an unbiased analysis of groups of genes within each brain region that are functionally related, co-occur and are statistically enriched in a given gene population. MOET uses the Gene Ontology (GO) consortium, in order to identify functional classes of genes that are statistically over-represented amongst the genes that are differentially expressed during HF (Smith *et al.*, 2020). For this analysis, we focused on the PFC, DH and CeA, which are the three brain regions that showed the most significant changes in gene expression and are implicated in the measures of mood and memory used in our experiments (see Fig.7; results from the PVN and VH are shown in Supplementary Fig.1. Note that MOET was not applied to BLA given that only 1 gene changed in this region). We highlight two major groups of GOs related to (1) biological, cellular and/or molecular pathways/functions (left graphs, 7A, C and E) and (2) disease-related processes (right graphs, 7B, D and F). For simplicity and relevance, only the GO subcategories in which the gene counts represented 50% or more of the total genes associated with that GO that were down- or upregulated during HF, and that were considered statistically significant by MOET are displayed.

**Figure 7.**
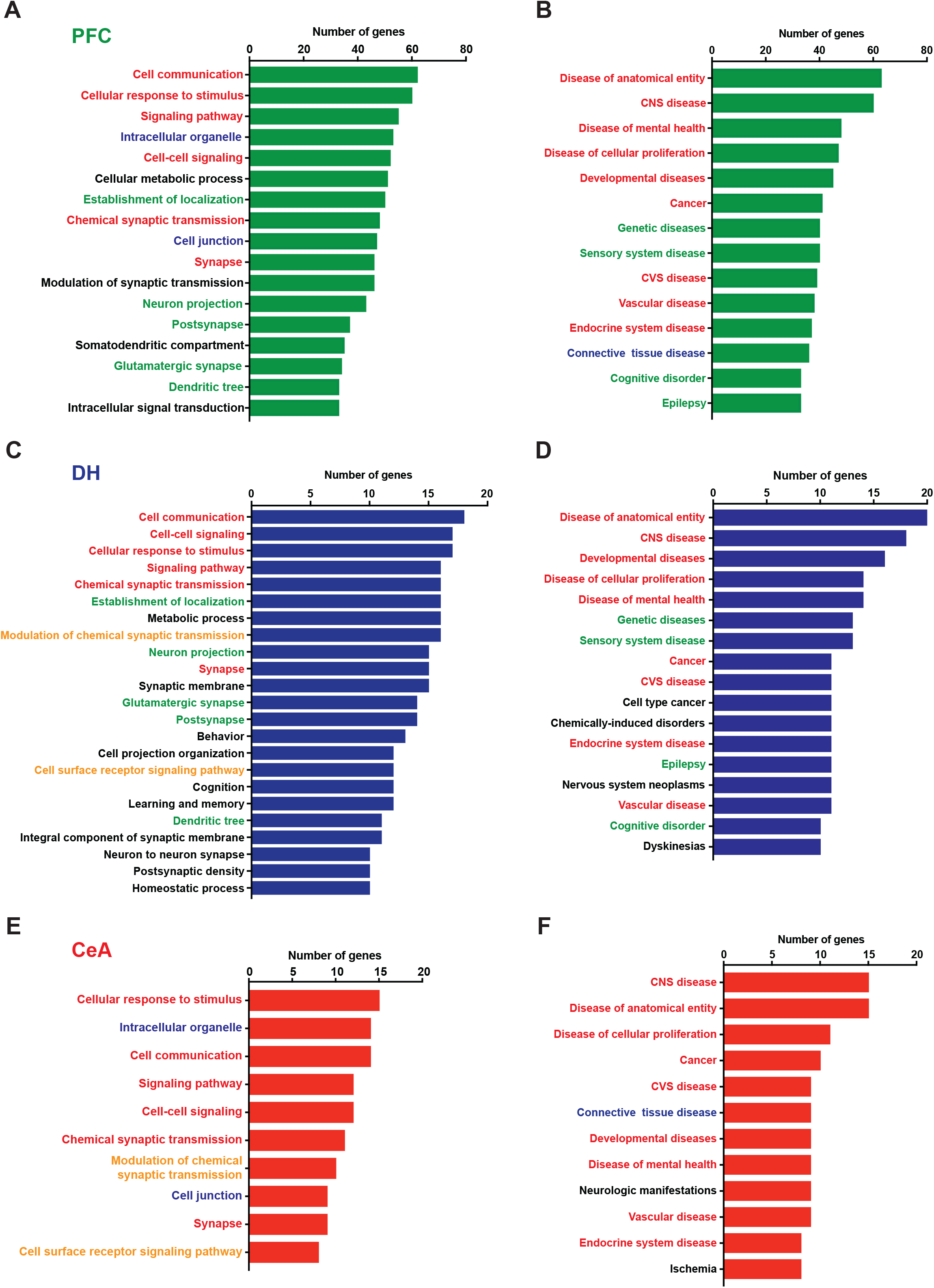
Quantitative ontology analysis of gene expression in the PFC, DH and CeA of HF rats. (**A,C,E**), Summary bar graphs showing the analysis of biological, cellular or molecular gene ontologies (GOs) terms that are significantly enriched based on the number of genes that were up- or downregulated in HF rats vs. Sham in PFC (**A**), DH (**C**) and CeA (**E**). (**B,D,F**) Summary bar graphs showing the analysis of disease ontology terms highlighting important disease categories that involve the up- or downregulated genes by HF in PFC (**B**), DH (**D**) and CeA (**F**). Categories written in red represent those enriched terms that show up in PFC, DH and CeA; green represents those that show up in PFC and DH; blue represents those that show up in PFC and CeA; and orange represents those that show up in DH and CeA. All the categories included in the graphs reached statistical significance p<0.00001.

For the first major group of GOs (Fig. 7A, C and E), the MOET analysis revealed that multiple genes that were robustly downregulated during HF are involved in cell communication, cell-cell signaling, cellular response to stimulus and synaptic structure/function/modulation, particularly for glutamatergic synapses (Fig. 7A and C, written in green). In addition, there was a notable involvement of genes involved in cellular metabolic processes and intracellular organelles. Interestingly, an involvement for genes related to learning and memory and cognition was observed to occur only in the DH (Fig. 7C). For the GOs related to disease processes (Fig. 7B, D and F), there was an enriched involvement of multiple genes across the three brain regions, which are associated with CNS diseases, diseases of mental health, as well as cognitive disorders. Importantly, the three regions also displayed involvement of genes involved in cardiovascular/vascular diseases. Finally, and perhaps somewhat surprising, an involvement of genes related to cancer/neoplasms was also observed.

In order to independently validate our PCR array data, we performed qRT-PCR using individual primers for five genes, *Adcy1*, *Arc*, *Homer1*, *Mapk1* and *Nr4a1,* with *B2m* as the housekeeping gene (see Fig. 8). In all cases, the qRT-PCR replicated the direction of the PCR array results (i.e., increase, decrease or no effects), although the magnitude did not match precisely. For example, the results with the individual primers showed that HF significantly downregulated *Adcy1* in the PFC (−1.30), PVN (−1.54), CeA (−1.49), DH (−1.48), and BLA (−1.12), but upregulated it in the VH (1.52), (Fig. 8A), which is the same pattern that we observed in the PCR array (Fig. 8B). A similar profile of results was observed for the individual primers and array for Arc, *Homer1*, *Mapk1* and *Nr4a1* (Fig. 8A and B).

**Figure 8.**
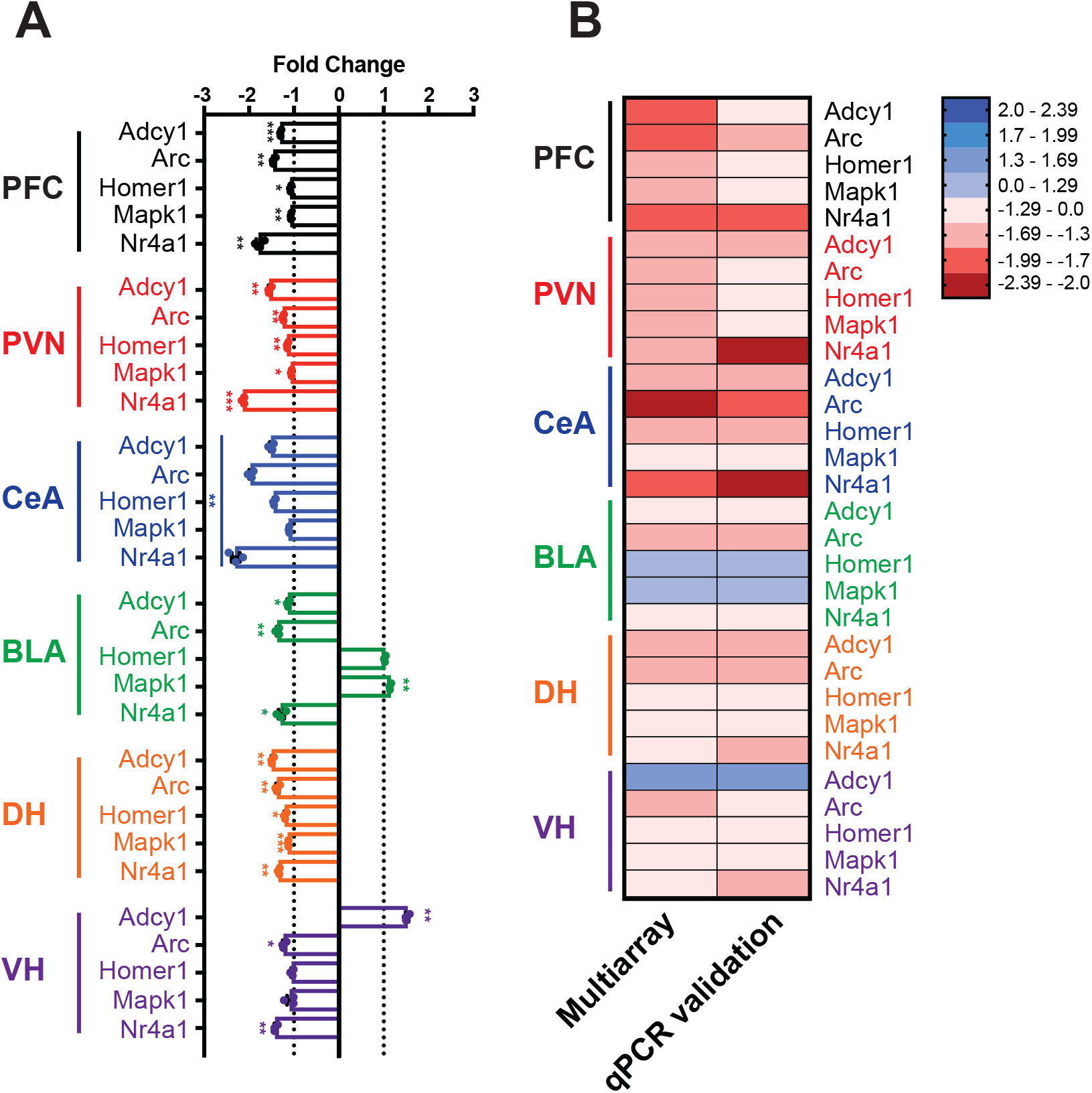
qRT-PCR validation of results obtained in the gene array study. (**A)**, Summary bar graphs showing the mean fold change in mRNA levels for *Adcy1*, *Arc*, *Homer1*, *Mapk1* and *Nr4a1* genes in HF rats in relation to Sham rats (theoretical mean = +/−1) in the PFC (black bars), PVN (red bars), CeA (blue bars), BLA (green bars), DH (orange bars) and VH (purple bars). Dashed lines indicate the theoretical means. (**B)**, heatmap graph showing the magnitude of change in gene expression in the array and validation shown as fold change in gene expression from −2.39 to +2.39 induced by HF in relation to sham rats’ gene expression. *p<0.05, **p<0.01 and ***p<0.001 vs. theoretical mean, one sample t-test.

## DISCUSSION

Collectively, the present findings show that a rat model of HF characterized by severely reduced ejection fraction disrupts several behavioral measures of mood and memory and drastically alters the pattern of synaptic plasticity-associated gene expression in brain areas critical for mood and memory. Specifically, we found that HF decreased sucrose preference, which is a sign of anhedonia, and increased measures of anxiety in the EPM. HF also impaired SA and IA retention performance, indicating that HF disrupts spatial working memory and emotional, long-term memory, respectively.

Although both clinical and experimental studies in animal models support a link between HF and cognitive/mood impairments, (Rutledge *et al.*, 2006; Angermann *et al.*, 2012; Lossnitzer *et al.*, 2013; Cameron *et al.*, 2014; Cannon *et al.*, 2017; Toledo *et al.*, 2019; Wang *et al.*, 2019), the precise mechanisms and molecular underpinnings remain elusive. Cognitive impairment in neurodegenerative disorders and aging, as well as mood disorders have been linked to altered synapse structure/functional properties as well as to deficits in synaptic plasticity mechanisms including blunted LTP. Thus, in order to gain insights into possible molecular underpinnings of cognitive and mood disorders in HF, we conducted a PCR array study that assessed the simultaneous expression of 84 genes critical for synaptic plasticity. Importantly, our findings showed that HF drastically downregulated numerous genes implicated in synaptic plasticity in a brain region-dependent manner, with the most profound changes observed in PVN and PFC, moderate changes in DH and CeA, and minimal alterations in VH and BLA.

To the best of our knowledge, the present results are the first to show that HF impairs SA and IA. Both memory tasks are dependent on the integrity of DH (Quevedo *et al.*, 1997; Izquierdo *et al.*, 2000; Dickson *et al.*, 2007; Sakaguchi & Sakurai, 2020) and PFC (Delatour & Gisquet-Verrier, 1996; Giovannini *et al.*, 2005; Torkaman-Boutorabi *et al.*, 2015; Woloszynowska-Fraser *et al.*, 2017), which are interconnected brain regions critical for memory. IA is also dependent on an intact VH, CeA, and BLA (Parent *et al.*, 1992; Roozendaal & McGaugh, 1997; Giovannini *et al.*, 2005; Nobre, 2013), which may account for the finding that HF-induced memory loss appeared more pronounced in the IA task than in SA. Collectively, the findings show that our animal model of HF impacts both working memory (*i.e*., SA) and long-term memory (i.e., IA), which is consistent with findings showing that rats with severely reduced ejection fraction have impaired long-term memory in the spatial water maze (Lu *et al.*, 2020) and with results showing that HF patients have deficits in both immediate and delayed recall (Leto & Feola, 2014). Short-term memory deficits appear to be one of the most pervasive cognitive issues in HF, which is significant because these problems are associated with impaired ability for HF patients to manage their medical condition (Gaviria *et al.*, 2011; Leto & Feola, 2014).

The present findings are consistent with previous research showing that HF decreases sucrose preference and increases anxiety in the EPM (Prickaerts *et al.*, 1996; Lu *et al.*, 2017; Najjar *et al.*, 2018b; Liu *et al.*, 2019; Wang *et al.*, 2019). The present results extend those findings by showing that these effects are observed in a rat model of HF characterized by severely reduced ejection fraction. Moreover, our gene array findings suggest that these effects are likely due to PFC dysfunction, given that manipulations that inhibit PFC produce anhedonia (Banasr & Duman, 2008; Kabir *et al.*, 2017) and increase anxiety (de Visser *et al.*, 2011).

Both the amygdala and the PVN are important neuronal substrates implicated in the regulation of mood, and recent studies support that a direct connection from the PVN to the amygdala modulates anxiety levels and fear memories (Knobloch *et al.*, 2012; Hasan *et al.*, 2019). Moreover, changes in proper neuronal function within the amygdala could contribute to the anxiogenic effect observed in HF rats (Lisboa *et al.*, 2010; Tye *et al.*, 2011; Strauss *et al.*, 2013; Ventura-Silva *et al.*, 2013; Sorregotti *et al.*, 2018; Pi *et al.*, 2020). Still, it is yet unclear whether the changes in gene expression reported in this study in the CeA, BLA and PVN could be directly related to the mood disorders reported in HF rats.

Our findings indicate that HF increases water intake, which is in agreement with a previous report indicating that myocardial infarction in rats and mice increase thirst (De Smet *et al.*, 2003; Frey *et al.*, 2014) and with evidence showing that excessive thirst is a common symptom observed in patients with HF (Allida *et al.*, 2015). This increase in thirst is likely due to chronic activation of the systemic and central angiotensin II signaling (Prager *et al.*, 2016) and/or increases in circulating levels of arginine vasopressin (Kheirbek *et al.*, 2012) that are both hallmarks of HF. Importantly, the PVN is a critical component of the neural circuitry that underlies thirst (Cohn *et al.*, 1984; Krishnan & Nestler, 2011; Dardiotis *et al.*, 2012; Dobryakova *et al.*, 2017) and our PCR gene array findings indicate that HF induces a massive decrease in gene expression in PVN.

Our findings show a drastic downregulation of multiple genes implicated in synaptic plasticity in a brain region-dependent manner in rats with HF compared to sham controls. For instance, changes in genes involved in learning and memory were observed only in DH and not in CeA or PFC. This may reflect the different role of these brain areas in cognition. DH involvement in SA and IA is assumed to reflect an episodic (*i.e.,* autobiographical) memory of where the animal has been (i.e., SA) and the context in which it received footshock (i.e., IA). In contrast, the amygdala’s role in IA may be more related to the emotional component of the task (Roozendaal *et al.*, 2009; Janak & Tye, 2015; Bocchio *et al.*, 2017) and PFC may be necessary for SA and IA due to its critical role in executive function and decision-making (Goldman-Rakic, 1996). Although spatial abilities and episodic memory appear to be the most compromised cognitive functions in HF (Hjelm *et al.*, 2012), HF patients do also show impaired executive function (Pressler *et al.*, 2010b; Harkness *et al.*, 2011; Dardiotis *et al.*, 2012).

The present results appear to be the first to show that HF impacts amygdala-dependent memory (i.e., IA) and significantly decreases the expression of several genes critical for synaptic plasticity in CeA, while minimally impacting another region of the amygdala implicated in mood and memory (i.e., BLA). To date, there is very limited evidence implicating the amygdala in any effects of HF in human patients or animal models. In patients, HF is associated with reduced blood flow to amygdala (Roy *et al.*, 2017) and increased amygdala responses to sympathetic activation (Woo *et al.*, 2005; Etkin & Wager, 2007). In animal models, HF increases glial density (GFAP immunoreactivity) in the medial amygdala, which is critical for cardiovascular reflexes and central control of sympathetic responses (Salazar *et al.*, 2014). HF also decreases orexin type 2 receptor gene expression in amygdala, although whether this decrease is restricted to a particular amygdala nucleus is unknown (Hayward *et al.*, 2015). Thus, the present results significantly advance our understanding of the consequences of HF by showing that HF characterized by a severe reduction in ejection fraction is associated with the downregulation of several genes in CeA and a relative sparing of BLA.

Our findings show that HF impacts DH expression but has minimal effect on VH, which is consistent with extensive evidence indicating that the hippocampus is not a unitary structure and that there are clear distinctions between DH and VH. For example, DH and VH have different anatomical connections, cellular and circuit properties and patterns of gene expression that likely contribute to the different functions that they serve (Moser & Moser, 1998; Thompson *et al.*, 2008; Dong *et al.*, 2009; Barkus *et al.*, 2010; Fanselow & Dong, 2010; Bienkowski *et al.*, 2018). DH appears to be primarily responsible for the cognitive functions of the hippocampus (Moser & Moser, 1998; Sahay & Hen, 2007; Barkus *et al.*, 2010; Fanselow & Dong, 2010), which is consistent with our finding that that genes associated with learning and memory were only impacted in DH. Our findings also add to evidence suggesting that DH is more vulnerable to neuroinflammation and other pathology than VH (Fuster-Matanzo *et al.*, 2011; Stouffer *et al.*, 2015; Dobryakova *et al.*, 2017; Dobryakova *et al.*, 2019; Tournier *et al.*, 2019).

Several critical families of genes were considerably affected in these brain regions in HF rats. These include various IEGs, such as *Arc*, *Egr*, *Jun* (but not *Fos*), which would suggest an overall reduction in neuronal activation in these areas, which could be a result of reduced inputs from upstream brain regions. Another major group of genes downregulated during HF were genes encoding glutamate receptors, predominantly metabotropic, but also ionotropic AMPA and NMDA receptor subunits. Given the importance of glutamate receptor signaling in maintaining appropriate basal network activity levels, as well as a playing a critical role in both up- and downregulation of the strength of synaptic efficacy (*e.g.,* LTP and LTD) (Park *et al.*, 2008; Waung *et al.*, 2008; Crupi *et al.*, 2019), it would be expected that the reported changes in GLURs may also contribute to lower levels of neuronal activation and compromised synaptic plasticity in HF rats. Other major gene families downregulated during HF included neurotrophic factors (e.g., BDNF, *Ntrk2* and *Ngf*) and kinases and phosphatases (e.g., *Ppp1cc*, *Ppp2ca*, *Prkca*, *Prkcg* and *Prkg1)*.

We used MOET to identify functional categories among the genes altered in HF within specific brain regions, which provides an unbiased analysis of groups of genes that are functionally related and that co-occurr and/or are co-affected during a specific condition (e.g., HF). For simplicity, we focused on two major Gene Ontology categories (GOs): (1) biological/molecular pathways and (2) disease-related processes. As summarized in Fig.7, the MOET analysis revealed that the majority of genes downregulated during HF were functionally related and are involved in cell communication, cellular response to stimulus, dendritic signaling, and synaptic structure/function/modulation, particularly for glutamatergic synapse. More importantly, MOET also revealed that multiple genes that changed across brain regions were associated with diseases of mental health, cognitive disorders as well as cardiovascular/vascular diseases. This is in agreement with multiple previous publications supporting that downregulation of several of the same genes we found in HF have been linked to cognitive dysfunction and neurodegenerative diseases such as Alzheimer’s Disease (Wong *et al.*, 1999; Guzowski *et al.*, 2000; Wang *et al.*, 2004; Plath *et al.*, 2006; Park *et al.*, 2008; Waung *et al.*, 2008; Davis *et al.*, 2010; Velazquez *et al.*, 2016; Wilkerson *et al.*, 2018).

Numerous putative mechanisms, not mutually exclusive, could contribute to the disruption of the molecular underpinnings of synaptic plasticity during HF. One obvious candidate is neuroinflammation, a hallmark pathophysiological phenomenon in HF (Diaz *et al.*, 2020). Activation of neuroinflammatory pathways within the hypothalamic PVN has been directly implicated as an underlying mechanism contributing to sympathohumoral activation in this disease (Kang *et al.*, 2008; Xu & Li, 2015; Yu *et al.*, 2018). Importantly, findings in rodents have implicated neuroinflammation and the expression of inflammatory and signal transduction genes in HF-induced cognitive impairment, depression and anxiety-like behavior (Frey *et al.*, 2014; Hay *et al.*, 2017; Toledo *et al.*, 2019; Wang *et al.*, 2019). This possibility is supported by evidence showing that HF impacts the expression of genes related to inflammation in the hippocampus and PFC of mice (Frey *et al.*, 2014).

Another critical pathophysiological mechanism in HF is cerebral hypoperfusion associated with low cardiac output that chronically leads to brain ischemia and microinfarcts (Celutkiene *et al.*, 2016). In fact, several clinical studies implicate compromised cerebral blood flow as a mechanism contributing to cognitive impairment and dementia in patients with HF (Pullicino & Hart, 2001; Zuccala *et al.*, 2001; Iadecola, 2013). It is noteworthy that hypoxia reduces the expression of genes that are required for the maintenance of synaptic structure and function (Bazan *et al.*, 2002), and several of the key genes found to be disrupted in HF in this study, including *Arc*, *Egr* and mGLURs were previously reported to be affected by hypoxia (Simonyi *et al.*, 1999; Hong *et al.*, 2003; Martin *et al.*, 2012; Zhao *et al.*, 2012; Alagappan *et al.*, 2013; Chan *et al.*, 2016; Chen *et al.*, 2016; Su *et al.*, 2020). Of particular interest to this study, *Arc* expression in the hippocampus was shown to be significantly downregulated following ischemic conditions, both *in vitro* and *in vivo* (Hong *et al.*, 2003). Moreover, manipulations that increase Arc expression protected against hypoxia-related cognitive impairment and hippocampal damage (Zeng *et al.*, 2019). Whether chronic brain hypoperfusion and/or neuroinflammation contribute to the disruption of molecular underpinnings of synaptic plasticity and cognitive and mood disorders in HF remains to be determined.

An important limitation of our studies is that they were focused exclusively on male rats. We chose to use males given the fact that most previous studies assessing the contribution of the hypothalamus to neurohumoral activation in the HF rat model, including our own (Biancardi *et al.*, 2011; Potapenko *et al.*, 2011; Potapenko *et al.*, 2012, 2013; Stern & Potapenko, 2013; Ferreira-Neto *et al.*, 2017; Ferreira-Neto & Stern, 2019), as well as those focusing on HF-associated neuroinflammation (Kang *et al.*, 2008; Xu & Li, 2015; Yu *et al.*, 2018), were conducted in male rats. Nonetheless, given that the incidence of HF and cognitive/mood comorbidities are sex-dependent (Gottlieb *et al.*, 2004; Rutledge *et al.*, 2006; Najjar *et al.*, 2018a), it is plausible that the behavioral and molecular changes reported here are sex-dependent as well.

Collectively, the present findings suggest that the impaired mood and cognition observed in HF patients that is associated with higher hospital readmission and mortality rates (Dickson *et al.*, 2007; Pressler *et al.*, 2010a; Lee *et al.*, 2013; Lossnitzer *et al.*, 2013) is likely due be due to pathology in DH and PFC. This interpretation is in accordance with the finding that learning and working memory deficits, delayed recall, depression, and anxiety are correlated with significant cerebral grey matter loss in hippocampus and PFC (Almeida *et al.*, 2012; O’Donnell *et al.*, 2012). Moreover, our findings suggest that the amygdala, particularly CeA, also contributes to mood and memory problems in HF. The present findings provide support for the fact that HF is not only a neurohumoral cardiovascular problem but is also a disorder of mood and memory. Moreover, our data also indicate that HF is associated with a drastic downregulation of several critical genes implicated in synaptic plasticity in brain areas necessary for cognitive, mood and homeostatic processes, suggesting that they could underlie the effects of HF on mood and memory.

## ADDITIONAL INFORMATION

### AUTHOR CONTRIBUTIONS

ARK conducted behavioral testing, data analyses and drafted portions of the manuscript; SK and KEW conducted behavioral testing and scoring and contributed to the literature review; HCFN performed HF surgeries, echocardiographic studies, brain punches, RNA extraction, array and qRT-PCR experiments and analysis, drafted the manuscript; FA performed brain punches for the array and qRT-PCR experiments; AAP and DNC contributed to the design, analysis, interpretation, figure construction and initial draft of the gene array and qRT-PCR data; MBP and JES designed the study, supervised experiments and drafted and edited the manuscript. All authors critically reviewed the manuscript and approved the final version for publication.

---

**Supplementary Figure 1.**
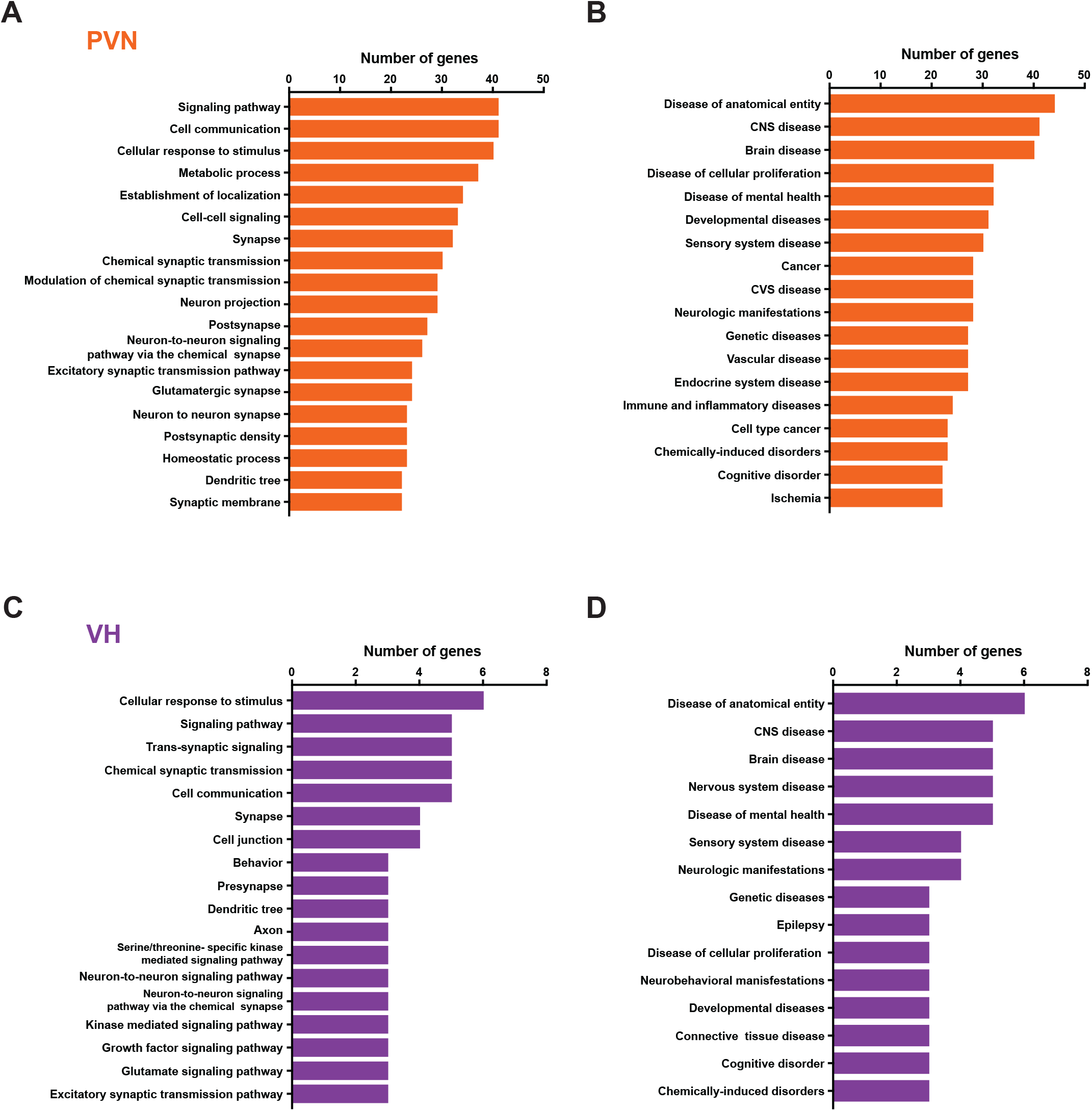
Qualitative ontology analysis of gene expression in the PVN and VH of HF rats. **(A)** and **(C)**, summary bar graphs showing the analysis of biological, cellular or molecular ontologies (GO) displaying critical biological processes categories that involves the up- or downregulated genes by HF in PVN **(A)** and DH **(C)**. **(B)** and **(D)**, summary bar graphs showing the analysis of disease ontology (GO) highlighting important disease categories that involves the up- or downregulated genes by HF in PVN **(B)** and DH **(D)**. All the categories in included in the graphs reached statistical significance p<0.00001, test used.

